# Surforama: interactive exploration of volumetric data by leveraging 3D surfaces

**DOI:** 10.1101/2024.05.30.596601

**Authors:** Kevin A. Yamauchi, Lorenz Lamm, Lorenzo Gaifas, Ricardo D. Righetto, Daniil Litvinov, Benjamin D. Engel, Kyle Harrington

## Abstract

**Motivation:** Visualization and annotation of segmented surfaces is of paramount importance for studying membrane proteins in their native cellular environment by cryogenic electron tomography (cryo-ET). Yet, analyzing membrane proteins and their organization is challenging due to their small sizes and the need to consider local context constrained to the membrane surface.

**Results:** To interactively visualize, annotate, and analyze proteins in cellular context from cryo-ET data, we have developed Surforama, a Python package and napari plugin. For interactive visualization of membrane proteins in tomograms, Surforama renders the local densities projected on the surface of the segmentations. Suforama additionally provides tools to annotate and analyze particles on the membrane surfaces. Finally, for compatibility with other tools in the cryo-ET analysis ecosystem, results can be exported as RELION-formatted STAR files. As a demonstration, we performed subtomogram averaging and neighborhood analysis of photosystem II proteins in thylakoid membranes from the green alga *Chlamydomonas reinhardtii.*

**Availability and implementation:** Python package, code and examples are available at: https://github.com/cellcanvas/surforama

## Introduction

Advances in cryogenic electron tomography (cryo-ET) provide the opportunity to view molecular structures in cellular context^1,2^. Localizing proteins in tomograms allows analyzing their 3D spatial organization and their interaction with native membranes. For example, cryo-ET has revealed the detailed spatial distribution of photosynthetic protein complexes in thylakoid membranes^3^, the geometry of COPII assembly on cargo vesicles^4^, the clustering of proteasomes at the ER membrane^5^, and the 3D organization of actin networks at the interface with the plasma membrane^6^. Accurate protein locations are an indispensable prerequisite to analyze these spatial relationships or perform subtomogram averaging to achieve high-resolution protein structures. However, this is a challenging task due to the low signal-to-noise ratio and anisotropic resolution (missing wedge) of cryo-ET data, as well as the necessity to consider the spatial relationships and cellular context in 3D. Small, membrane-embedded proteins are particularly difficult to identify and annotate because large portions of the proteins are embedded within the membrane, and the geometric information of the membrane’s surface needs to be considered. Here, we describe Surforama, an open-source software tool for visualizing and annotating membrane proteins by leveraging surface geometries, enabling human-in-the-loop curation of particle annotations for large-scale automated pipelines.

## Prior art

While many tools have been developed for the visualization and analysis of membranes in tomograms^7,8^, few offer the ability to view and annotate data on surfaces. Membranorama^3^ provides essential functionalities in this regard, but it is a standalone application, limiting its integration into workflows. Membranorama cannot integrate additional data (e.g., segmentations of other cellular structures) into the visualization and is limited to volumes that fit in GPU memory. Further, it is only available on the Windows operating system.

ChimeraX^9^, a popular visualization tool in structural biology, is capable of rendering volumetric data directly on surfaces such as those obtained by membrane segmentation from tomograms. Combined with the ArtiaX plugin^10^, it is possible to pick and orient particles on the segmentations within ChimeraX. While its rendering capabilities are extremely powerful, ChimeraX lacks specialized tools for membrane analysis such as controlling the thickness of the projection on the segmentation surface and constraining the rotation of particles oriented normal to the membrane. Similarly, the blik^11^ napari^12^ plugin offers several capabilities for annotation of particles in tomograms, including the consideration of different membrane geometries for pre-orienting the particles and straightening of segmentations for visualization purposes. However, it also lacks features dedicated to the inspection of membrane surfaces.

To provide a performant, interactive tool for visualizing, annotating, and analyzing membranes in cryo-ET data, we have developed Surforama (https://github.com/cellcanvas/surforama), an open source Python library. This tool allows users to pick particles on membranes and explore cryo-ET tomograms by projecting tomographic densities along surfaces. By building on top of napari^12^ and the scientific Python ecosystem, Surforama is easy to extend and combine with other tools.

All of these approaches, including Surforama, rely on the availability of accurate membrane segmentations. Recent advances in computational methods^13,14^ have facilitated the generation of such membrane segmentations in cryo-ET data.

### Features

Surforama is a Python package for visualizing, annotating, and analyzing the organization of particles on a surface, with the target application of studying the molecular organization of membranes imaged by cryo-ET (Fig. 1A). Surforama functionality can be accessed via the core Python library and a graphical user interface (GUI) for interactive analysis. The GUI is accessible both via a command line interface and as a napari plugin (Fig. 1B).

**Figure 1.**
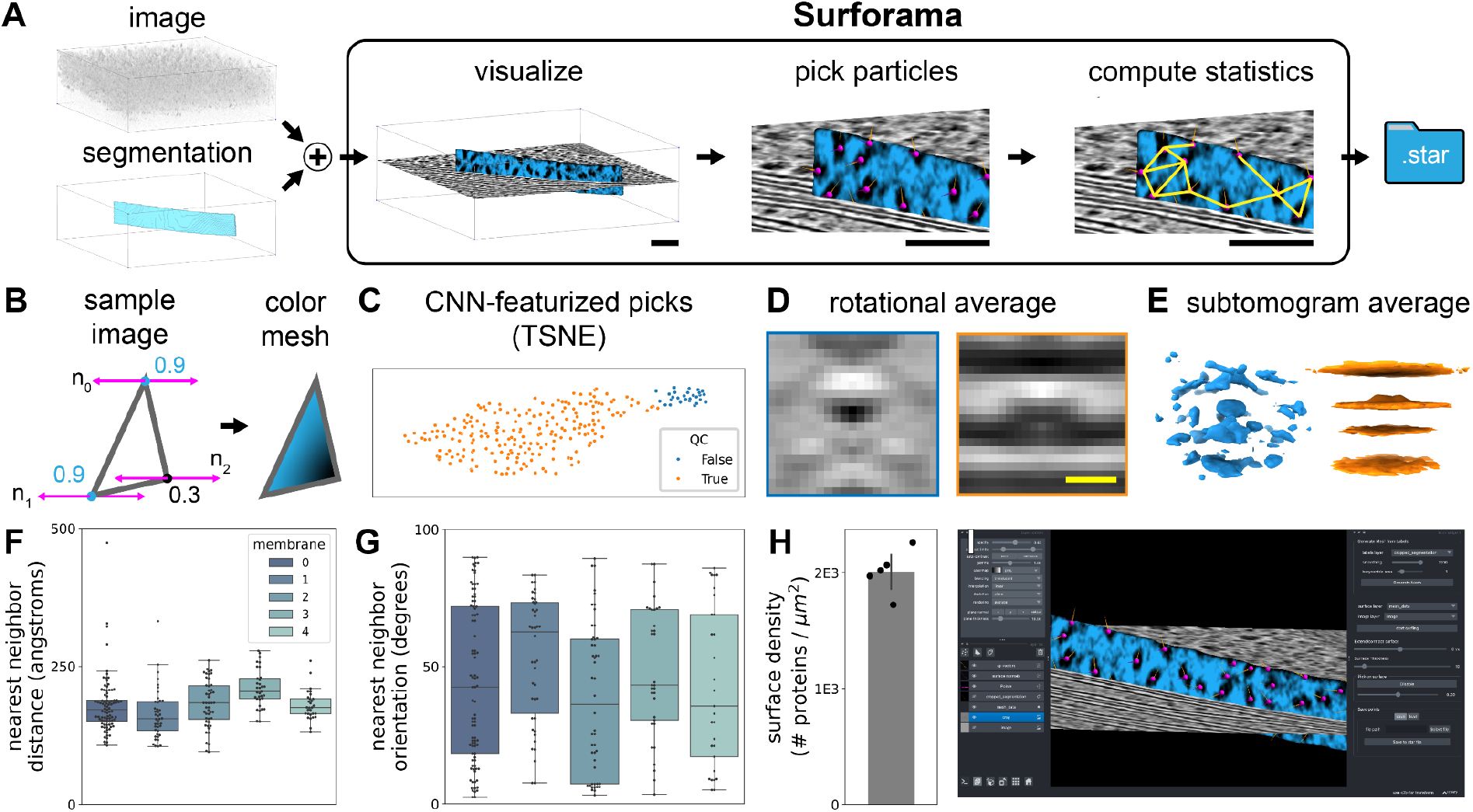
Surforama is an interactive application for human-in-the-loop particle picking and analysis on membrane surfaces. **(A)** Schematic of Surforama workflow demonstrated with cryo-ET imaging of thylakoid membranes inside Chlamydomonas cells, where photosystem II protein complexes were picked (magenta spheres) and their neighborhood graph constructed by geodesic distance along the surface (yellow lines). **(B)** Densities of proteins protruding from the membrane are projected onto the mesh surface by sampling the tomogram data along mesh vertex normals. The mean intensity is then assigned to the vertices and interpolated across the mesh element. **(C)** Demonstrating the utility of integrating with the scientific Python ecosystem, we used a pre-trained convolutional neural network to featurize 2D rotational averages extracted from the pick locations (2D t-SNE plot of the featurized picks shown). Using this featurization, we clustered the picks to automatically filter out those with erroneous orientations. **(D)** Representative 2D rotational averages of the particle locations assigned to each class (left: false positive, right: true positive). **(E)** 3D subtomogram averages calculated with STOPGAP from the false (left, blue) and true (right, orange) positive classes identified by our analysis of the photosystem II particles. Surforama provides functions to quantify the organization of molecules on the surface including the **(F)** nearest neighbor geodesic distances and **(G)** the orientation of particles relative to their neighbors. **(H)** Surface densities of proteins can be computed and compared across multiple membranes. **(I)** To make Surforama widely accessible and integrate visualization of data from other imaging modalities, we have implemented the graphical user interface as a napari plugin. Scale bars are 50 nm in and 5 nm in (D).

Surforama provides preprocessing functions to convert tomograms and membrane segmentations into surfaces upon which local densities can be visualized. In particular, Surforama converts the membrane segmentation from a dense voxel representation by first meshing with the marching cubes algorithm and then remeshing the surface using Voronoi clustering^15^. Surface normals are then computed for each element in the mesh.

To visualize the local densities on a membrane surface, the meshes are rendered in napari and colored by the densities sampled at the surface (Figure 1A). The densities are sampled at each vertex of the mesh by interpolating the intensity value at a point along the normal vector from the vertex position (Figure 1B). The length of the normal vector can be changed smoothly to visualize the membrane-adjacent context. Further, the mesh surface can be expanded and contracted to sample different distances from the segmented surface. This efficient implementation allows users to run Surforama without a powerful GPU or dedicated workstation.

Proteins can be picked directly in the napari interface by clicking locations on the mesh in the napari viewer (Figure 1A). Importantly, users can additionally assign protein orientations: The 3D orientation is specified by two vectors: the mesh normal vector and an orthogonal user-specified vector corresponding to the in-plane orientation. The particle poses are stored in a Pandas data frame and can be exported as a RELION-formatted STAR file^16^ to enable a seamless transition to subtomogram averaging and other downstream applications.

To analyze the molecular organization on the surface of the membrane, Surforama provides functions to construct and quantify neighborhood features. Neighbors are identified using geodesic distances. Once neighborhoods are constructed, one can measure nearest neighbor distances and compare the orientation of particles relative to the neighbors (Figure 1F,G). Surforama can also quantify membrane-level features such as the particle density (Figure 1H).

Surforama has been designed to be interoperable with the large ecosystem of scientific Python software as well as domain-specific tools. To enhance interoperability, Surforama uses standard in-memory objects (e.g., numpy arrays^17^). We ensure compatibility with cryo-ET tools for important tasks such as subtomogram averaging by outputting oriented particles in RELION-formatted star files^16^. The Surforama GUI is implemented as a napari plugin (Figure 1I), which allows integration with visualization of other data (e.g., wider cellular context from a tomogram). Surforama can be installed on Mac OS, Windows, and many versions of Linux.

## Results

As a case study, we used Surforama to analyze 5 thylakoid membranes from the green alga *Chlamydomonas reinhardtii* using data from a previous study^3^. The tomograms were originally segmented using TomoSegMemTV^18^, followed by cropping the membranes of interest in Amira (ThermoFisher Scientific). We converted these segmentations to triangular meshes using Surforama’s built-in functionalities. Using the surface picking tool, we picked 241 oriented locations centered on putative photosystem II proteins. Highlighting the advantage of integration with the scientific Python ecosystem, we then adapted PySeg’s 2D rotational averaging functionality^19^ to generate averages of the picked particles generated by Surforama (Fig 1C, D). We utilized a ResNet50 convolutional neural network pre-trained on ImageNet^20^, followed by k-means clustering, to perform automated quality control, sorting out picks where either position or orientation seem to be faulty. The clusters become evident by plotting the data along t-SNE feature vectors (Figure 1C). Compared to expert-annotated ground truth, this automated quality control performed well (100% recall, 80% precision for outlier detection). The two classes were further validated by calculating 3D subtomogram averages in STOPGAP^21^, using the particle coordinates and orientations manually assigned when picking in Surforama (Fig. 1D, E), without further refinement. To assess how the photosystem II proteins are organized on the membranes, we quantified the orientation of the particles relative to their neighbors (Fig 1F). Finally, we compared the density of proteins in different membranes (Fig. 1G).

## Conclusion

Surforama is an open-source Python library for visualizing cryo-ET images of membranes and annotating particle locations for downstream analysis (e.g., subtomogram averaging). We have designed Surforama such that it matches existing community standards, allowing it to be integrated with other tools. While we have initially targeted cryo-ET, the integration with the napari ecosystem facilitates use in other domains of bioimaging (e.g., volume EM and 3D fluorescence microscopy). To ease adoption, we have provided online documentation with tutorials on key use cases. We anticipate that Surforama will enable a new frontier for the analysis of membrane-bound structures.

## Code availability

Surforama can be installed via pip. The source code is available at https://github.com/cellcanvas/surforama under an MIT license. Installation and usage instructions are available at https://github.com/cellcanvas/surforama.

## Conflicts of interest

None declared.

## Funding

This work was supported by the Biozentrum of the University of Basel, the Chan Zuckerberg Imaging Institute, the Munich School for Data Science (MUDS), a Boehringer Ingelheim Fonds fellowship to L.L., and an ERC consolidator grant “cryOcean” (fulfilled by the Swiss State Secretariat for Education, Research and Innovation, M822.00045) to B.D.E..

## Notes

### Competing Interest Statement

The authors have declared no competing interest.

## References

1. Navarro, P. P. Quantitative Cryo-Electron Tomography. Front. Mol. Biosci. 9, 934465 (2022).

2. Young, L. N. & Villa, E. Bringing Structure to Cell Biology with Cryo-Electron Tomography. Annu. Rev. Biophys. 52, 573–595 (2023).

3. Wietrzynski, W. et al. Charting the native architecture of thylakoid membranes with single-molecule precision. Elife 9, (2020).

4. Pyle, E. & Zanetti, G. Cryo-electron tomography reveals how COPII assembles on cargo-containing membranes. bioRxiv 2024.01.17.576008 (2024) doi:10.1101/2024.01.17.576008.

5. Albert, S. et al. Direct visualization of degradation microcompartments at the ER membrane. Proceedings of the National Academy of Sciences 117, 1069–1080 (2020).

6. Jasnin, M. et al. Elasticity of podosome actin networks produces nanonewton protrusive forces. Nat. Commun. 13, 1–11 (2022).

7. Barad, B. A., Medina, M., Fuentes, D., Wiseman, R. L. & Grotjahn, D. A. Quantifying organellar ultrastructure in cryo-electron tomography using a surface morphometrics pipeline. J. Cell Biol. 222, (2023).

8. Salfer, M., Collado, J. F., Baumeister, W., Fernández-Busnadiego, R. & Martínez-Sánchez, A. Reliable estimation of membrane curvature for cryo-electron tomography. PLoS Comput. Biol. 16, e1007962 (2020).

9. Meng, E. C. et al. UCSF ChimeraX: Tools for structure building and analysis. Protein Sci. 32, e4792 (2023).

10. Ermel, U. H., Arghittu, S. M. & Frangakis, A. S. ArtiaX: An electron tomography toolbox for the interactive handling of sub-tomograms in UCSF ChimeraX. Protein Sci. 31, e4472 (2022).

11. Gaifas, L., Timmins, J. & Gutsche, I. blik: an extensible napari plugin for cryo-ET data visualisation, annotation and analysis. bioRxiv 2023.12.05.570263 (2023) doi:10.1101/2023.12.05.570263.

12. napari: a multi-dimensional image viewer for Python. doi:10.5281/zenodo.8115575.

13. Lamm, L. et al. MemBrain v2: an end-to-end tool for the analysis of membranes in cryo-electron tomography. bioRxiv 2024.01.05.574336 (2024) doi:10.1101/2024.01.05.574336.

14. Moreno, J. J., Garzón, E. M., Fernández, J. J. & Martínez-Sánchez, A. HPC enables efficient 3D membrane segmentation in electron tomography. J. Supercomput. 78, 19097–19113 (2022).

15. GitHub - pyvista/pyacvd: Python implementation of surface mesh resampling algorithm ACVD. GitHub https://github.com/pyvista/pyacvd.

16. Burt, A. et al. An image processing pipeline for electron cryo-tomography in RELION-5. bioRxiv 2024.04.26.591129 (2024) doi:10.1101/2024.04.26.591129.

17. Harris, C. R. et al. Array programming with NumPy. Nature 585, 357–362 (2020).

18. Robust membrane detection based on tensor voting for electron tomography. J. Struct. Biol. 186, 49–61 (2014).

19. Martinez-Sanchez, A. et al. Template-free detection and classification of membrane-bound complexes in cryo-electron tomograms. Nat. Methods 17, 209–216 (2020).

20. Deng, J. et al. ImageNet: A large-scale hierarchical image database. 10.1109/CVPR.2009.5206848.

21. Wan, W., Khavnekar, S. & Wagner, J. STOPGAP: an open-source package for template matching, subtomogram alignment and classification. Acta Crystallographica Section D: Structural Biology 80, 336–349 (2024).

